# Cyclin C promoter occupancy directs changes in stress-dependent transcription

**DOI:** 10.1101/2020.07.14.202580

**Authors:** David C. Stieg, Katrina F. Cooper, Randy Strich

## Abstract

The Cdk8 kinase module (CKM) is a detachable Mediator subunit composed of cyclin C, and one each of paralogs Cdk8/Cdk19, Med12/Med12L and Med13/Med13L. In addition to regulating transcription, a portion of cyclin C also leaves the nucleus following cytotoxic stress to induce mitochondrial fragmentation and apoptosis. Our previous RNA-seq studies demonstrated that cyclin C represses a subset of hydrogen peroxide-induced genes under normal conditions, while also being required for the full induction of other loci following stress. Here, we show that cyclin C directs this transcriptional reprograming through changes in its promoter occupancy. Following peroxide stress, cyclin C promoter occupancy increased for genes it activates while decreasing at loci it represses under normal conditions. Promoter occupancy of other CKM components generally mirrored cyclin C indicating that the CKM moves as a single unit. However, CKM integrity appeared compromised at a subset of repressed promoters suggesting a source of cyclin C that is targeted for nuclear release. Interestingly, mTOR inhibition induced a new pattern of cyclin C promoter occupancy indicating that this control is fine-tuned to the individual stress. Using inhibitors, we found that Cdk8 kinase activity is not required for CKM movement or repression but was necessary for full gene activation. In conclusion, this study revealed that different stress stimuli elicit specific changes in CKM promoter occupancy correlating to altered transcriptional outputs. Finally, although CKM components were recruited or expelled from promoters as a unit, heterogeneity was observed at individual promoters suggesting a mechanism to generate gene- and stress-specific responses.

## Introduction

Cells exhibit multiple adaptive responses following exposure to cytotoxic agents. The failure to mitigate the cellular damage caused by these stressors can initiate regulated cell death (RCD) pathways (1). One common cause of cellular damage is reactive oxygen that can be derived either endogenously (e.g., elevated respiration, oxidase overexpression) or by environmental exposure to pro-oxidants (2) such as H_2_O_2_ or chemotherapeutics (3). An important early adaptation is transcriptional reprogramming, which directs many molecular processes within the cell. The transcriptional adaptations induced by oxidative stress include the upregulation of both pro-death and prosurvival genes. For example, oxidative stress stimulates the tumor suppressor p53-dependent activation of genes that promote intrinsic RCD (iRCD) (4). In addition, genes encoding proteins mitigating the effects of oxidative stress are also induced (4). This system allows the cell to balance the amount of damage incurred with its ability to facilitate repairs. In contrast to oxidative stress, amino acid starvation inhibits the mammalian target of rapamycin (mTOR) resulting in induction of several genes necessary for cell survival pathways including autophagy (5). However, prolonged mTOR inhibition represses cell growth eventually inducing cell death (6). Therefore, the balance between survival and death is influenced by the type, intensity and duration of the stress that is relayed to the nucleus to impact transcription.

Our previous RNA sequencing (RNA-seq) studies identified the oxidative stress cyclin C-regulated transcriptome in mouse (*Mus Musculus*) embryonic fibroblast (MEF) cells (7). Cyclin C was originally identified as a transcriptional repressor of meiotic and stress response genes in the yeast *S. cerevisiae* (8–11). However, mammalian cyclin C functions as both a transcriptional activator and repressor, with an essential role in development (12–14). Cyclin C is a component of the Cdk8 Kinase Module (CKM), which is a detachable and transiently interacting subunit of the Mediator complex (15). The other members of the CKM include Cdk8 or Cdk19, Med12 or Med12L, and Med13 or Med13L (16). Cdk8/Cdk19, Med12/Med12L and Med13/Med13L are paralogs with similar but not identical functions that are incorporated independently into CKMs, along with cyclin C and Cdk8, in a 1:1:1:1 stoichiometry (17). Given this dual role in transcription, it may not be surprising that genetic and epidemiological studies have provided evidence that the CKM can either stimulate or suppress tumorigenesis (18). For example, *CDK8* overexpression stimulates colon and breast cancer progression (19,20). Accordingly, Cdk8 inhibitors have been developed and tested in preclinical studies demonstrating a positive impact within multidrug combination regimens (21). Conversely, deleting *Ccnc* (cyclin C) stimulates hyperplasia in thyroid and acute lymphoblastic leukemia mouse models (18,22). Interestingly, there is evidence that the CKM regulates transcription in both a CDK8 kinase-dependent and kinase-independent manner (23). These studies indicate that the CKM plays a complicated role in transcriptional control responding to diverse inputs. However, how the CKM itself is controlled to mediate this dual transcriptional activity is unclear.

In addition to its nuclear role, ~15-20% of cyclin C, but not Cdk8, translocates to the cytoplasm in response to oxidative stress (24). In addition to inactivating Cdk8 to alter transcription, cytoplasmic cyclin C directly binds both the dynamin-like GTPase Drp1 and the pro-apoptotic factor Bax to promote mitochondrial fragmentation and efficient iRCD execution, respectively (24–26). This mitochondrial role is conserved in yeast and our laboratory demonstrated that Med13 is required for cyclin C nuclear retention (27). To allow cyclin C nuclear release in response to oxidative stress in yeast, Med13 is degraded via the ubiquitin proteasome system (UPS) in a highly regulated process requiring multiple signaling pathways and Cdk8 activity (28,29). Similarly, structural studies revealed that Med13/Med13L tether the CKM to the Mediator complex suggesting that its role in retaining cyclin C in the nucleus is conserved (30). However, it is not clear how the pool of cyclin C is selected for nuclear release in mammalian cells and whether Med13/Med13L destruction is involved. Here, chromatin immunoprecipitation (ChIP) studies revealed that promoter recruitment or loss of cyclin C and the CKM directly correlated with its role as a transcriptional activator or repressor. This profile changed following starvation stress indicating that cyclin C and CKM movements are precisely determined by the insult. Finally, CKM integrity was disrupted at a subset of repressed promoters suggesting a source for cyclin C nuclear release. Taken together, these findings are consistent with a model that the CKM is a dynamic complex that directs transcriptional repression and activation through changes in promoter occupancy at target genes.

## Results

### Cyclin C transcriptional regulation of oxidative stress-responsive genes

Our transcriptome analysis revealed that cyclin C mediates both positive and negative transcriptional control of genes induced by H_2_O_2_ exposure (7). To begin investigating this observation, we used gene ontology (GO) (31) to identify biological processes enriched in the data set for genes induced by H_2_O_2_ treatment. We chose to study two pathways, p53-induced genes and those required for autophagy as they represent stresses leading to cell death and survival, respectively. The p53 stress response genes included *Trp53inp1* (Tumor protein p53-inducible nuclear protein 1), *p21* (*Cdkn1a*; Cyclin-dependent kinase inhibitor 1) and *Jmy* (Junction-mediating and -regulatory protein) (labeled in figures in ***bold italic font***). The autophagic genes chosen were *Gabarap* (Gamma-aminobutyric acid receptor-associated protein) and *Prkaa2* (5’-AMP-activated protein kinase catalytic subunit alpha-2) (labeled in figures in <u>*underlined italic font*</u>). In addition, three genes *Adrm1* (Proteasomal ubiquitin receptor Adrm1), *Gtf2h1* (General transcription factor IIH subunit 1) and *Crebrf* (CREB3 regulatory factor) were chosen that were induced by H_2_O_2_ but are neither the p53-dependent nor autophagy responsive genes (labeled in figures in *italic font).* Finally, *Sall1* (Sal-like protein 1) and *Col4a1* (Collagen alpha-1(IV) chain) were chosen as cyclin C repressed and activated their transcription under non-stress conditions, respectively but their transcript levels are not affected by H_2_O_2_ stress.

We first confirmed the transcriptional changes observed with RNA-seq by RT-qPCR analysis. As expected, the three gene sets exhibited various levels of induction in response to H_2_O_2_ treatment while *Sall1* and *Col4a1* were not affected (Fig. 1A). Two patterns of transcription were observed for cyclin C-activated genes. For the p53-dependent set, cyclin C is required for steady state levels of *Trp53inp1* and *p21* but accounted for only partial H_2_O_2_ induction (left side, Fig. 1A). The transcriptional analysis of nine additional H_2_O_2_-induced genes controlled by p53 identified seven that required cyclin C for activation and two that were repressed during normal growth (Fig. S1). The same patterns were observed in that cyclin C is required for maintaining both steady state transcription and full H_2_O_2_-induced induction. A second pattern was observed for *Adrm1* and *Gabarap* transcription in that cyclin C was required for both steady state levels and any H_2_O_2_-mediated induction. For the cyclin C-repressed genes, two transcription patterns were again observed depending on p53 requirement. Deleting *Ccnc* resulted in a fourfold derepression of *Jmy* mRNA that was not enhanced by H_2_O_2_ treatment (right side, Fig. 1A). However, H_2_O_2_-treated WT cells only displayed a twofold elevation in *Jmy* mRNA suggesting the pro-oxidant treatment only partially relieves cyclin C-dependent repression. Similar to *Jmy* regulation, two repressed genes (*Dyrk3* and *Noxa*) exhibited elevated mRNA levels in the absence of cyclin C beyond that induced by H_2_O_2_ treatment in wild-type cells (Fig. S1). These findings suggest that H_2_O_2_ treatment does not elicit full activation of these loci. A second profile was observed for autophagy induced *Prkaa2* and two additional repressed genes *Gtf2h1* and *Crebrf*. These loci displayed mRNA levels in Ccnc-/- that matched WT H_2_O_2_ treated cells (Right side, Fig. 1A). Moreover, the addition of H_2_O_2_ did not increase transcription in *Ccnc^-/-^* cells. These findings suggest that H_2_O_2_ treatment was able to remove all cyclin C-dependent repression.

**Figure 1:**
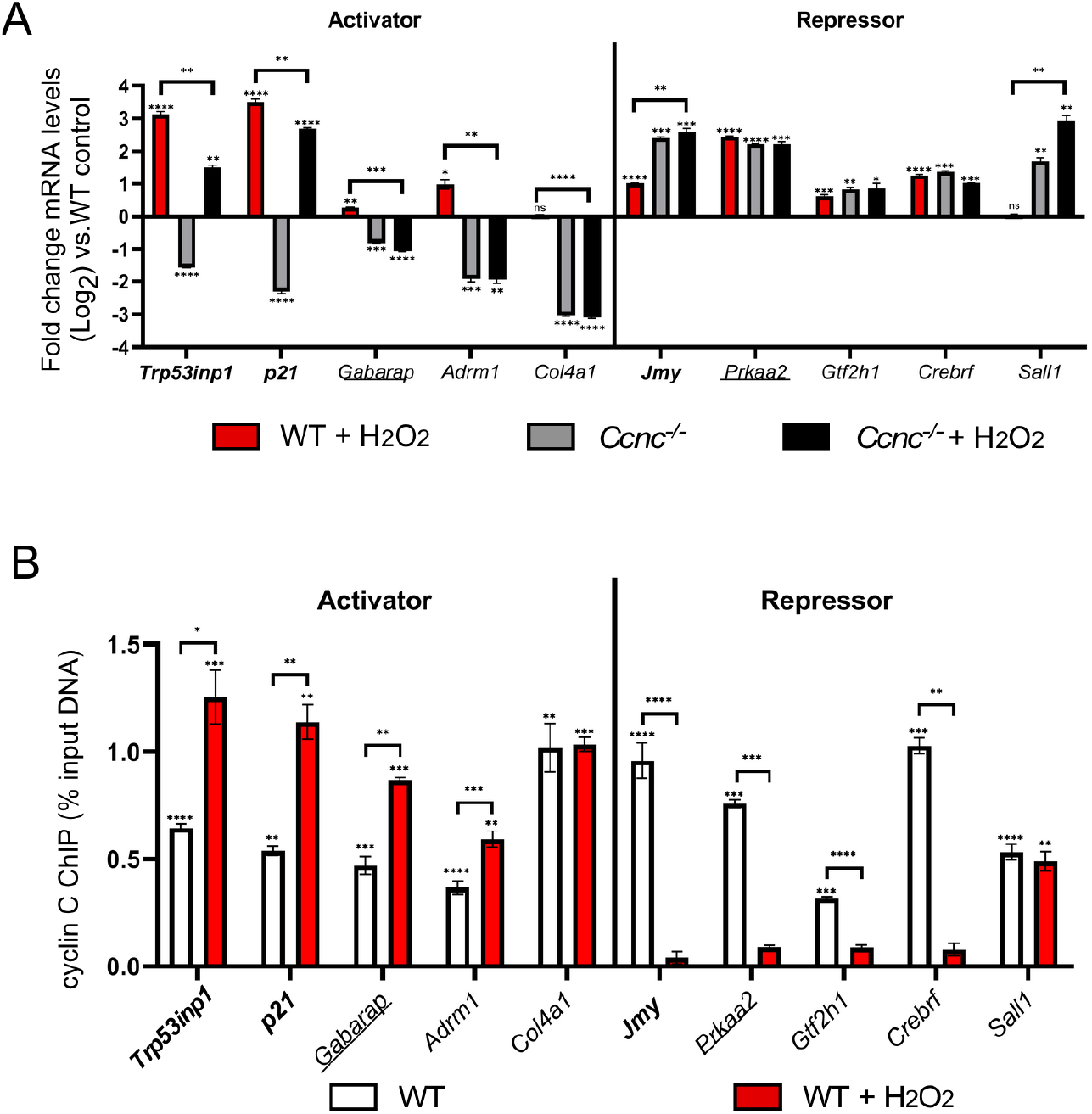
Cyclin C transcriptional regulation of oxidative stress-responsive genes. (A) RT-qPCR analysis for the genes indicated in WT or *Ccnc^-/-^* MEF cultures before or after H_2_O_2_ treatment as indicated. Results are given as fold change from WT untreated control. Values are presented ± WT untreated control (?og2). (B) ChIP analysis was quantified by qPCR for target promoter DNA that was first normalized to the input DNA control then compared to a non-specific antibody control (GFP). Values are presented as % input for a standard fraction of chromatin solution. Genes indicated in bold face are members of the p53 regulon. Genes indicated by underlining are also induced by autophagic conditions. Remaining loci represent other genes induced by H_2_O_2_ but are not members of the first two sets. For all results, statistical significance is indicated by the following: * p-value < 0.05, ** p-value < 0.01, *** p-value < 0.001, **** p-value < 0.0001.

Next, ChIP analysis was used to monitor cyclin C promoter occupancy at these promoters before and after H_2_O_2_ treatment. For all genes, cyclin C demonstrated promoter enrichment over background in unstressed cells (Fig. 1B) suggesting that its control of transcription at each locus is direct. Antibody specificity controls are shown in Fig. S2 for the *Trp53inp1* locus. Following H_2_O_2_ exposure, there was a significant increase in promoter occupancy for the two p53-dependent genes activated by cyclin C (*Trp53inp1* and *p21*) (left side, Fig. 1B). Previous reports demonstrated increased promoter occupancy at the *p21* promoter by the CKM members cyclin C, Cdk8 and Med12 following Nutlin, but not UVC, treatment (32). In contrast, there was a significant decrease in promoter occupancy of cyclin C following oxidative stress for the repressed gene *Jmy* (right side, Fig. 1B). These results indicate that cyclin C is recruited to promoters when its transactivation ability is employed but removed when its repression function is no longer wanted. Similar increases in occupancy were observed for the activated autophagy *Gabarap* and independent *Adrm1* genes. Likewise, loss in cyclin C occupancy was observed for the repressed genes *Prkaa2, Gtf2h1*, and *Crebrf* consistent with removing CKM repression. No changes in promoter occupancy were observed for the H_2_O_2_ independent genes *Col4a1* and *Sall1.* Taken together, these results are consistent with a model that cyclin C recruitment helps direct normal gene induction while its removal is important for transcriptional derepression.

### The promoter occupancy of CKM components largely mirrors that of cyclin C

Our results indicate that cyclin C occupancy is increased when it serves a positive role in transcription but removed to relieve its repressor function. Biochemical studies purifying the CKM suggest that it is mostly found as an intact complex (15). Therefore, we next asked whether other components of the CKM also display similar dynamic changes in promoter occupancy. ChIP experiments were conducted using antibodies targeting three additional members of the CKM (Cdk8, Med13 and Med13L). Med13 and Med13L were chosen as their individual activities have not been clearly delineated. The results indicated significant promoter occupancy by each of the four members for each gene in WT unstressed cells for both cyclin C-activated (Fig. 2A) and repressed (Fig. 2B) genes. Following H_2_O_2_ exposure, promoter occupancy patterns of the CKM members generally mirrored that observed for cyclin C for each of the induced genes *Trp53inp1, p21, Gabarap*, and *Adrm1* (Fig. 2A). As expected, no change in promoter occupancy for any of these proteins was observed for H_2_O_2_-independent *Col4a1.* For the repressed genes *Jmy* and *Gtf2h1*, there was a significant decrease in promoter occupancy of each CKM member in response to oxidative stress similar to that observed for cyclin C (Fig. 2B). However, CKM promoter occupancy at *Prkaa2* and *Crebrf* gave an intermediate result. Specifically, although cyclin C levels were diminished to nearly background levels at these promoters, significant levels of the remaining CKM members were detected (arrows, Fig. 2B). These findings indicate that cyclin C was separated from other CKM components at a subset of the *Prkaa2* or *Crebrf* promoters. Interestingly, ChIP analysis showed that Cdk8 occupancy at the *Trp53inp1* promoter was at the limits of detection in *Ccnc^-/-^* cells (Fig. S2). These results suggest a different fate for Cdk8 promoter occupancy when cyclin C is completely absent versus removed in response to stress. Together, these data indicate that promoter movements by cyclin C and other CKM components are largely similar arguing that the CKM moves as a single unit to and from promoters. A potential caveat to this conclusion is the two repressed promoters, *Prkaa2* and *Crebrf*, where cyclin C and CKM removal appear disconnected.

**Figure 2:**
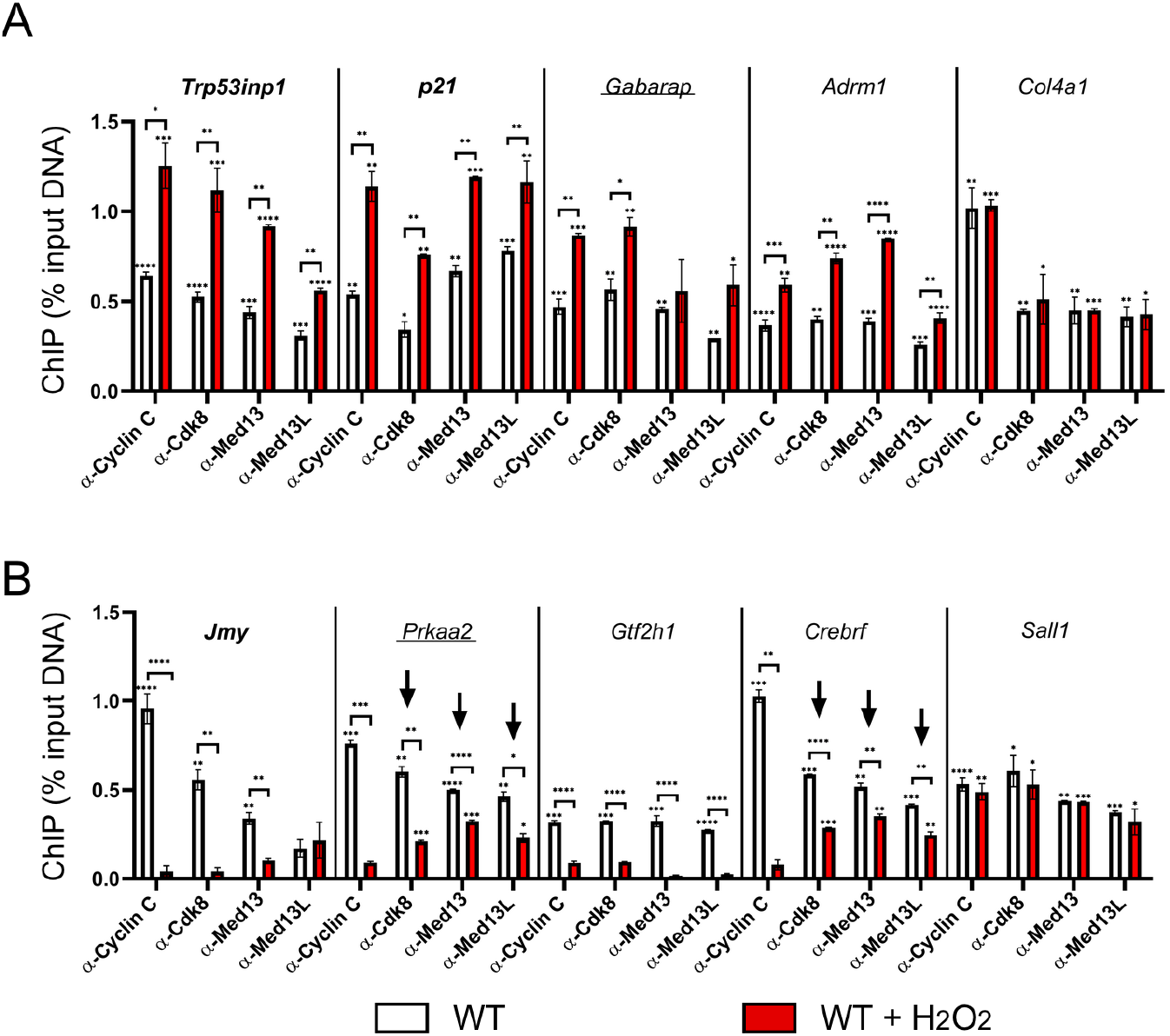
CKM members maintain similar but not identical promoter occupancy patterns at target genes. ChIP analysis for the CKM components indicated are shown cyclin C-activated (A) and -repressed (B) loci before and following H_2_O_2_ treatment. Two genes whose mRNA expression is not altered by H_2_O_2_ (*Col4al* and *Salll*) are shown. Arrows indicate genes displaying a disconnect between loss in cyclin C occupancy compared to the other CKM components. Data obtained from ChIP was compared to a non-specific control antibody (GFP). For all results, statistical significance is indicated by the following: * p-value < 0.05, ** p-value < 0.01, *** p-value < 0.001, **** p-value < 0.0001.

### A subset of genes are regulated by cyclin C in response to both oxidative stress and mTOR inhibition

Previous studies have revealed regulatory crosstalk between the oxidative stress and autophagic signaling systems (33). Specifically, it has been reported that low-level oxidative stress induces autophagy (34) while amino acid starvation triggers a low-level oxidative stress response (35). To determine if cyclin C exhibited coregulation of H_2_O_2_ and autophagy induced genes, WT MEF cultures were treated with the mTOR inhibitor Torin1 [32]. Similar to H_2_O_2_ stress, cyclin C was required for both steady state and Torin1-induced transcription of *Gabarap* and *Adrm1* (left side, Fig. 3A). In addition, two repressed genes, *Prkaa2* and *Crebrf* exhibited derepression following Torin1 treatment that was similar to those observed in *Ccnc^-/-^* cells, suggesting that removing cyclin C-dependent repression was sufficient for maximal induction (right side, Fig. 3A). Interestingly, one gene repressed by cyclin C, *Gtf2h1*, exhibited reduced mRNA levels following Torin1 treatment (arrow, Fig. 3A). These finding indicate that Torin1 and H_2_O_2_ treatment is processed similarly though cyclin C activity although differences exist.

**Figure 3.**
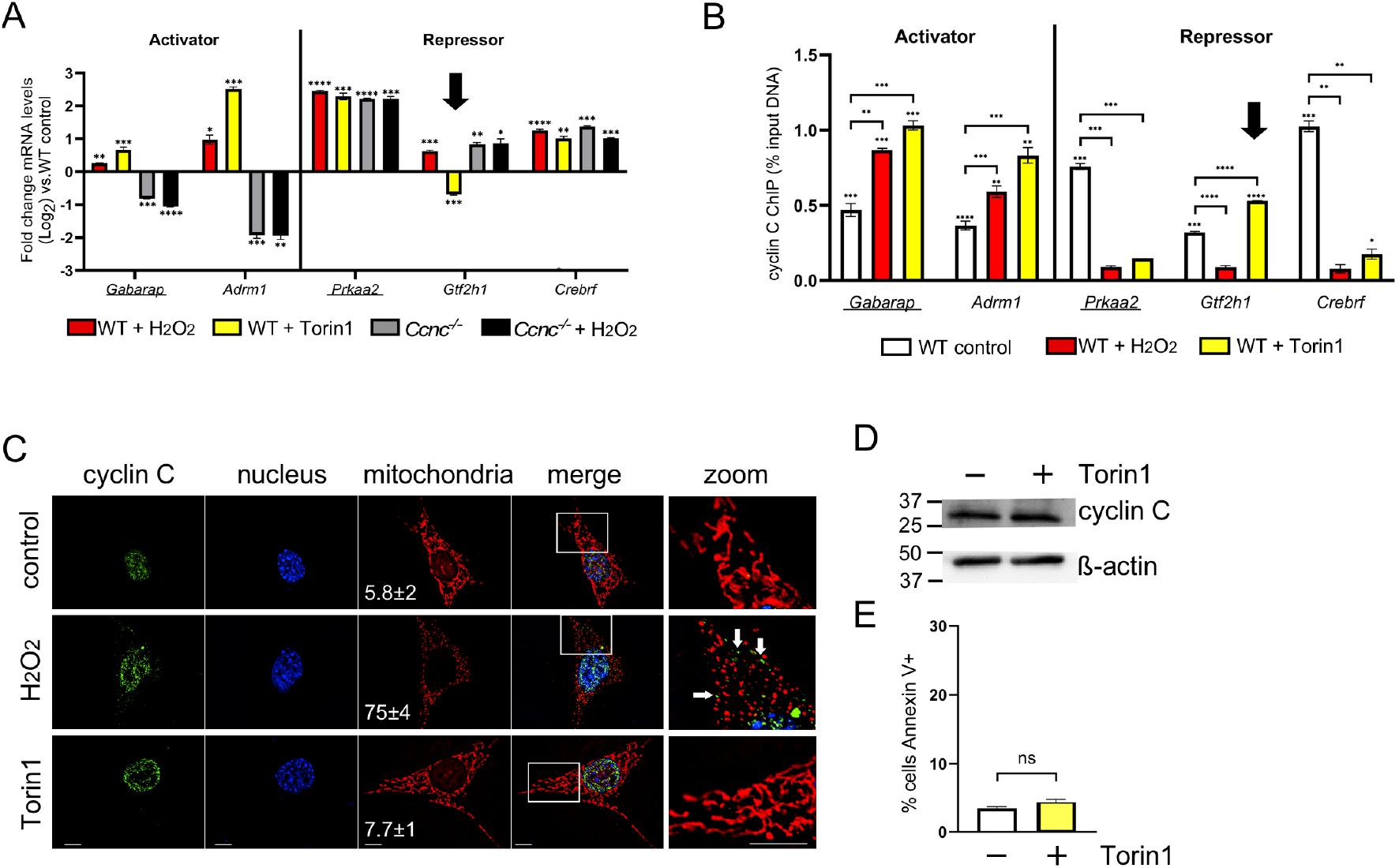
Cyclin C activates and represses an overlapping gene set induced following H_2_O_2_ and Torin1 treatment. (A) mRNA levels for the indicated genes were determined by RT-qPCR following exposure to oxidative stress and Torin1. Genes known to be upregulated under starvation conditions are underlined. Results from Fig. 1 for *Ccnc^-/-^* and WT cells are shown for comparison. Arrow indicates *Gtf2h1* repression following Torin 1 treatment. (B) cyclin C ChIP analysis of the genes described in (A). The arrow indicates the elevated occupancy of cyclin C at *Gtf2h1* co-incident with increased repression. (C) Subcellular localization of cyclin C was determined by immunocytochemistry. Mitochondria and nuclei were identified with MitoTracker Red and DAPI staining, respectively. The scale in the lower left corner of the images measures 10 nm, 40 nm for Zoom image. The percent of the cells exhibiting mitochondrial fragmentation is indicated (n = 3, ±S.D). (D) Cyclin C levels in cultures following exposure to Torin1 (250 nM, 4 h). β-actin was used as a loading control. Molecular weight markers (kDa) are indicated. (E) The percentage of the population Annexin V positive in WT cells following Torin1 treatment was determined by flow cytometry. Statistical significance is indicated by the following: * p-value < 0.05, ** p-value < 0.01, *** p-value < 0.001, **** p-value < 0.0001, ns, not significant.

Similar to H_2_O_2_ stress, ChIP analysis following Torin1 exposure revealed increased cyclin C promoter occupancy for the induced genes *Gabarap* and *Adrm1* (Fig. 3B, left side). In addition, cyclin C promoter occupancy was reduced at both *Prkaa2* and *Crebrf* promoters. These results are consistent with our model that removing cyclin C from promoters is part of the derepression process. Although the impact of cyclin C is opposite, these data suggest the possibility that a common pathway is used to induce these loci following both H_2_O_2_ exposure and mTOR inhibition. However, *Gtf2h1* mRNA levels decreased following Torin1 treatment (arrow, Fig. 3A), opposite to that observed for H_2_O_2_ stress. As our results indicate that cyclin C is recruited to promoters demonstrating transcriptional induction, we would predict that its promoter occupancy would also increase following Torin1 treatment to establish additional Torin1-induced repression. ChIP analysis proved this prediction correct with a significant increase in cyclin C promoter occupancy following Torin1 treatment compared to untreated cells (Fig. 3B). Further analysis of the p53-mediated stress response genes *Trp53inp1, p21* and *Jmy* found no or little change in mRNA levels between control and Torin1 treated cells (Fig. S3A). Consistent with this result, cyclin C promoter occupancy also remained unchanged (Fig. S3B). Our ChIP results from the repressed *Prkaa2* and *Crebrf* promoters suggested a possible source for nuclear released cyclin C following H_2_O_2_ treatment (see Fig. 2). Similarly, cyclin C is removed from these two promoters in response to Torin1 treatment. However, unlike H_2_O_2_ exposure, cyclin C remained nuclear following Torin1 treatment and the mitochondria remained intact (Fig. 3C). Finally, there was no significant difference in cyclin C levels (Fig. 3D) or iRCD induction (Fig. 3E) in response to Torin1 treatment. These results suggest that not only is CKM occupancy altered in response to a particular stimuli, its integrity may also be precisely controlled. These findings underscore the flexibility the cell has in employing cyclin C to fine tune transcription and mitochondrial responses to various stressors.

### Locus-specific release of cyclin C is regulated by the ubiquitin-proteasome system (UPS)

Under normal growth conditions, mammalian Med13 and Med13L exhibit a steady state turnover mediated by the UPS (36). This study found that Med13 or Med13L degradation correlated with decreased CKM-Mediator association (36). In yeast, Med13 degradation in response to oxidative stress is also mediated by the UPS (27). Therefore, we tested whether H_2_O_2_-induced cyclin C promoter release and transcriptional derepression required the UPS. First, wild-type cells were treated with the proteasome inhibitor MG-132 prior to H_2_O_2_ treatment. Analysis of the mRNA levels from the cyclin C repressed genes exhibited three different responses. The mRNA levels of the p53 regulated gene *Jmy* were unchanged compared to H_2_O_2_ treatment alone (Fig. 4A). Conversely*, Gtf2h1* and *Crebrf* transcription was elevated above H_2_O_2_ treatment alone suggesting that both stressors contributed to their increased transcription. Finally, H_2_O_2_-induced *Prkaa2* mRNA induction was reduced upon MG-132 addition. This result suggested that MG-132 inhibited full derepression. ChIP analysis for the genes *Jmy, Gtf2h1* and *Crebrf* showed a reduction in promoter occupancy following exposure to the combination treatment, comparable to that of oxidative stress alone (Fig. 4B). However, although cyclin C occupancy was reduced at the *Prkaa2* promoter following treatment with both drugs, the level of reduction was significantly less than observed in H_2_O_2_ treated cells. These results are consistent with a model that MG-132 prevented Med13 and/or Med13L proteolysis allowing at least partial retention of the CKM and reduced transcriptional repression. If correct, we would predict that Med13 and/or Med13L levels would be reduced in response to H_2_O_2_ but this effect blunted by UPS inhibition. Consistent with this possibility, Western blot analysis of Med13 and Med13L in extracts prepared from MEF cultures treated with H_2_O_2_ revealed an reduction of about 20% in protein levels compared to the untreated controls (Fig. 4C, quantified in Fig. 4D). However, this reduction was reversed upon addition of MG-132. Taken together, these results are consistent with the previous results from yeast and mammalian cells that cyclin C promoter release is associated with Med13/Med13L degradation. Importantly, not all promoters demonstrated elevated cyclin C retention following MG-132 treatment is consistent with our previous findings that the regulation of cyclin C occupancy, and CKM integrity, is controlled in a locus-specific manner.

**Figure 4:**
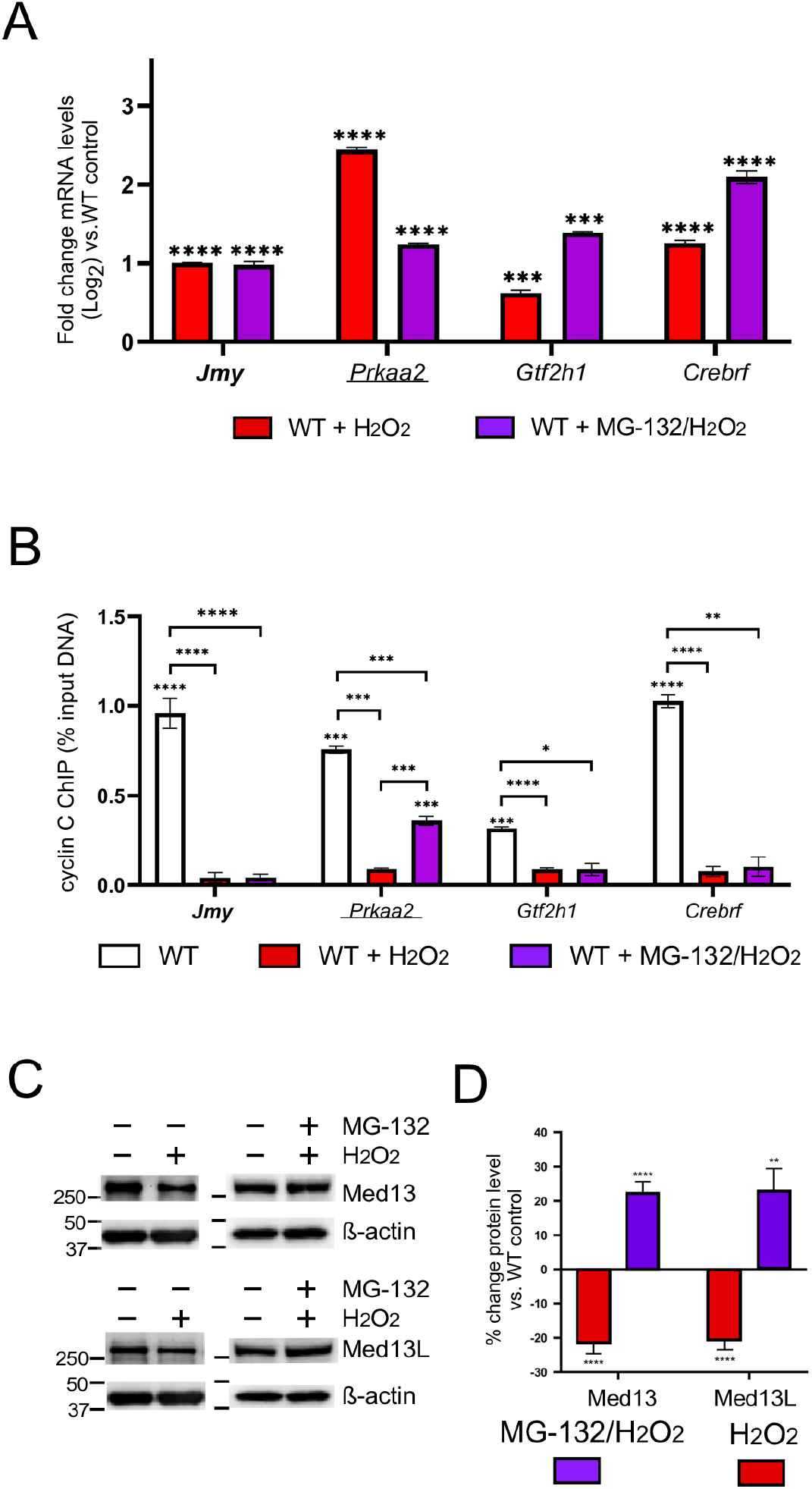
Release of cyclin C from promoters does not require Medl3/Medl3L degradation. RT-qPCR (A) and ChIP analysis (B) for the cyclin C repressed genes indicated following H_2_O_2_/MG-132 treatment. ChIP data are shown as percent input DNA compared to a non-specific control (GFP antibody). H_2_O_2_ treated WT control is from Fig. 1 to aid in comparing results. (C) Med13 and Med13L levels were monitored by Western blot analysis in MEFs following exposure to H_2_O_2_ and MG-132 as indicated, β-actin was used as the internal loading control. (D) The ratios of Med13 and Med13L signals to the β-actin internal control were calculated then compared WT untreated control (n=3). Statistical significance is indicated by the following: * p-value < 0.05, ** p-value < 0.01, *** p-value < 0.001, **** p-value < 0.0001.

### Cyclin C regulates transcription both dependent and independent of Cdk8 kinase activity

Our yeast studies revealed a Cdk8 phosphorylation is required for Med13 destruction and subsequent cyclin C nuclear release into the cytoplasmic (28). To test whether Cdk8 activity is required for the changes observed for the CKM promoter occupancy in mammalian cells, we directly compared transcription and promoter occupancy in *Ccnc^-/-^* mutants and WT cells were treated with the Cdk8 kinase inhibitor Senexin A (37). Following 24 h Senexin A treatment, mRNA levels of cyclin C-activated genes *Trp53inp1, p21, Gabarap*, and *Adrm1* were reduced comparable to the *Ccnc^-/-^* cells (left side, Fig. 5A). These data indicate that before stress, steady state transcription of these genes is dependent on Cdk8 kinase activity. Similarly, the p53-dependent genes *Trp53inp1* and *p21* were induced to levels similar to that observed in *Ccnc^-/-^* MEFs following H_2_O_2_ and Senexin A treatment (Fig. 5A). Moreover, the two additional cyclin C-activated genes, *Gabarap* and *Adrm1*, remained downregulated following Senexin A treatment phenocopying the *Ccnc*^-/-^ deletion results (Fig. 5A). These results indicate that Cdk8 kinase activity is required for both steady state transcription and induction following H_2_O_2_ treatment although for differing levels depending on the locus. The examination of cyclin C promoter occupancy for these activated genes revealed that cyclin C recruitment to these promoters following H_2_O_2_ treatment did not require Cdk8 kinase activity (left side, Fig. 5B).

**Figure 5.**
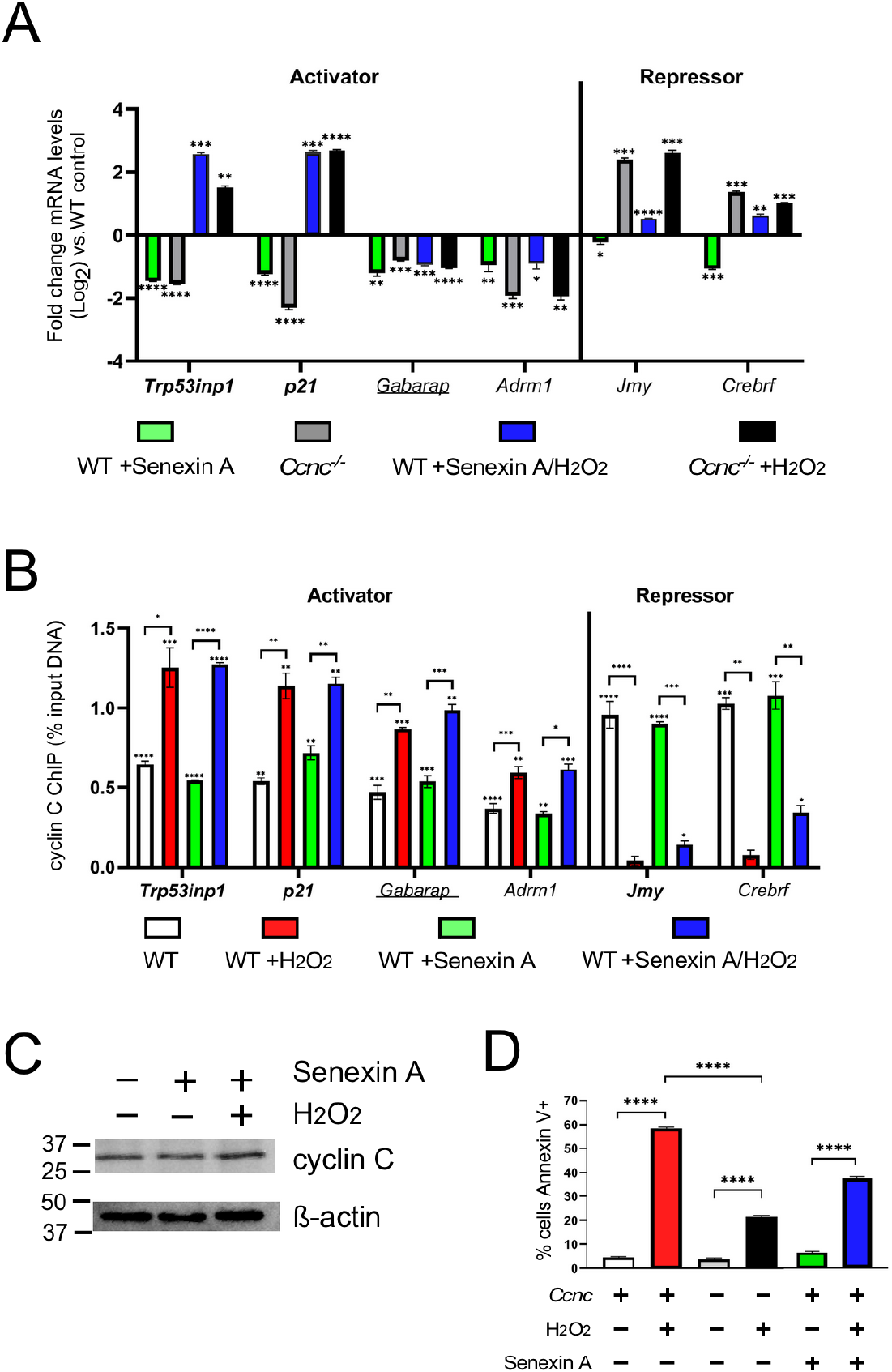
Cyclin C transcriptional activation, but not repression, is dependent on Cdk8 kinase activity. (A) RT-qPCR (A) and ChIP (B) analyses are shown for the indicated genes following treatment with Senexin A and H_2_O_2_. Values for H_2_O_2_-treated cultures are from Fig. 1 to aid in comparisons. (C) Cyclin C levels in MEFs following Senexin A treatment (1 μM, 24 h), and Senexin A + H_2_O_2_ (0.4 mM, 4 h) were determined by Western blot analysis, β-actin was used as a loading control. (D) The percentage of the population Annexin V positive following Torinl treatment was determined by flow cytometry (n=3). Statistical significance is indicated by the following: * p-value < 0.05, ** p-value < 0.01, *** p-value < 0.001, **** p-value < 0.0001.

Surprisingly, the two cyclin C repressed genes, *Jmy* and *Crebrf*, were slightly downregulated following Senexin A treatment compared to controls (right side, Fig. 5A). This is in contrast to *Ccnc^-/-^* MEFs that exhibited derepression of both genes. These results indicate that transcriptional repression of *Jmy* and *Crebrf* is independent of Cdk8 kinase activity. The upregulation of *Jmy* and *Crebrf* following exposure to oxidative stress is associated with the promoter release of cyclin C. In comparison to WT unstressed MEFs, ChIP analysis shows loss of cyclin C occupancy with Senexin A treatment (right side, Fig. 5B). However, subtle differences were observed with cyclin C release in Senexin A + H_2_O_2_ treated cells at the *Crebrf* promoter. Specifically, the gene *Crebrf* showed significantly more cyclin C at its promoter under these conditions (p-value = 0.02; Student’s T-test). However, the observed increase in cyclin C promoter occupancy for the gene *Jmy* was not statistically significant. These data indicate that although differences in cyclin C promoter occupancy are largely independent of Cdk8 kinase activity, its complete removal following H_2_O_2_ stress may be compromised. These differences are not due to changes in overall cyclin C levels following Senexin A addition (Fig. 5C). Interestingly, there was a significant decrease in iRCD efficiency in Senexin A treated cells following exposure to oxidative stress (Fig. 5D). These results indicate that Cdk8 kinase activity was required for efficient iRCD. This reduction was not as severe as that observed in *Ccnc*^-/-^ cells (Fig. 5D) suggesting differences exist between Senexin A treatment and *Ccnc* deletion for iRCD control. This difference may be due to the additional role of cyclin C at the mitochondria in oxidatively stressed cells (see Discussion). Taken together, these data demonstrate that Cdk8 kinase activity is required for full transcriptional activation while repression appears independent of this function.

## Discussion

The Cdk8 kinase module (CKM) interacts with the Mediator complex to induce the transcription of genes responsive to several stress signaling pathways including p53-mediated (32), oxidative (7) or hypoxia (38). In addition, we found that cyclin C represses steady state transcription of genes that are also induced by H_2_O_2_ treatment (7). However, mechanisms underlying how the CKM performs both positive and negative locus-specific roles in response to the same stress remain unclear. As the CKM is a removable, sub-stoichiometric component of Mediator, this report tested whether its gain or loss from promoters provided a regulatory mechanism to achieve this control. Using the CKM component cyclin C as a probe to monitor promoter occupancy, we examined both activated and repressed loci that respond to two stressors, H_2_O_2_-induced oxidative stress and starvation through mTOR inhibition. As observed in previous reports (32,38), we found that cyclin C is recruited to promoters that require the CKM for stress-induced activation. In addition, we found that cyclin C occupancy also increases when additional repression is required. Conversely, cyclin C is removed from stress-induced genes repressed by the CKM under normal conditions. These patterns of cyclin C occupancy are specific to whether H_2_O_2_ or starvation stress was administered suggesting that changes in CKM promoter occupancy may provide a strategy to achieve this complicated regulatory system. Our results also revealed that the other CKM members largely followed cyclin C promoter recruitment or release patterns, indicating that this complex moved as a unit although exceptions were noted (see below). Given the relatively low concentration of CKM in the cell, these results suggest that the CKM being released from a promoter undergoing derepression has two fates. First, the intact CKM is recruited to a promoter requiring this complex for transcriptional activation (Fig. 6). The other path involves dissolution of the CKM to allow cyclin C alone to translocate to the cytoplasm where it interacts with proteins at the mitochondria to induce fission and iRCD. Taken together, these results reveal that the CKM exhibits remarkable locus-specific movement to and from promoters based on its transcriptional role. This heterogeneity in response observed at individual promoters suggests that this system offers the cell a myriad of combinations with which to combine transcriptional control with CKM fate.

**Figure 6.**
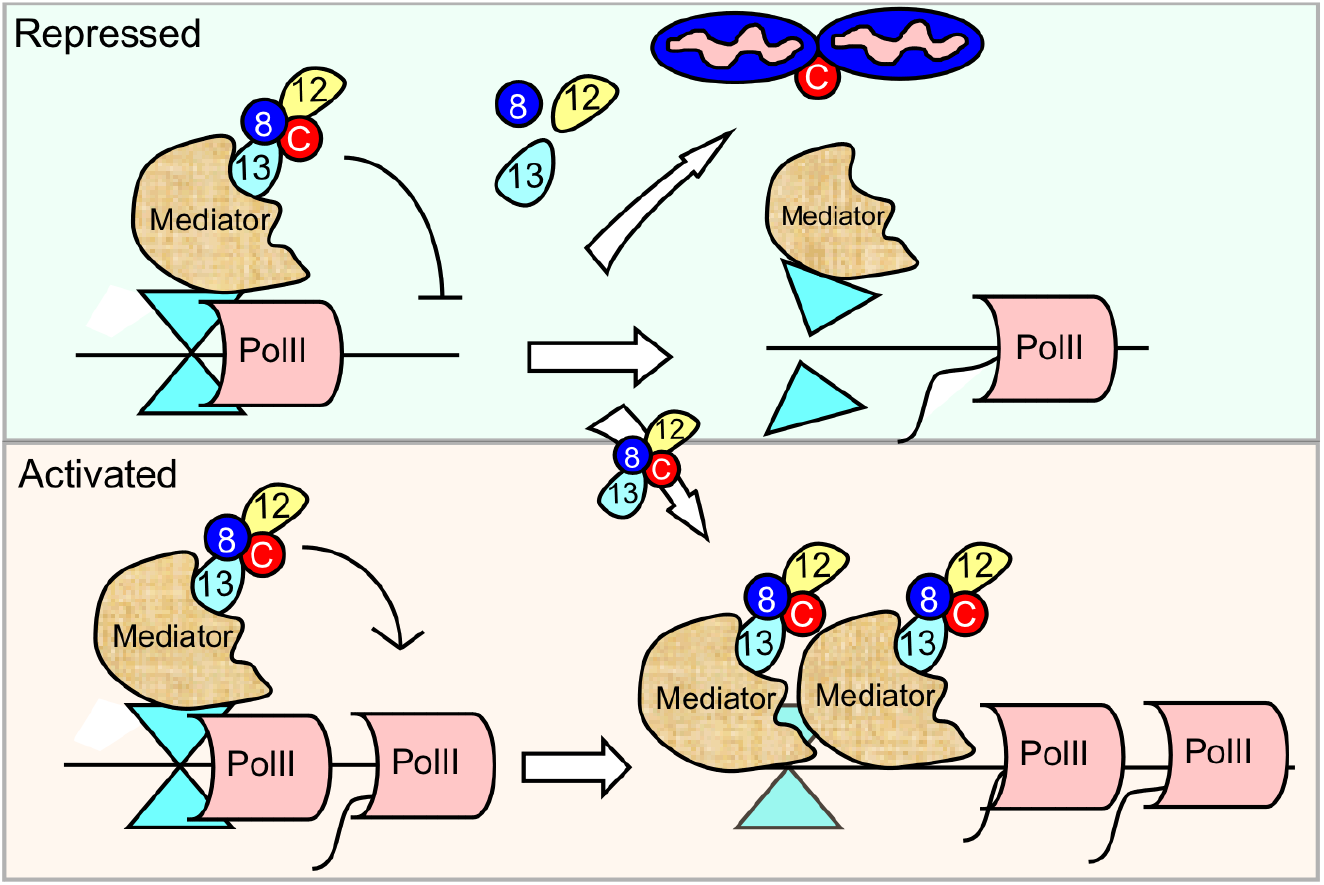
Model for CKM fate following H_2_O_2_ stress treatment. Examples of cyclin C repressed (top) and activated (bottom) genes are shown. The stress signal induces CKM release from genes repressed under normal conditions. The CKM has two fates. Intact CKM leaving the repressed promoters transits to loci to stimulate transcription (bottom panel). A subpopulation of CKMs dissolves allowing free cyclin C to exit the nucleus (top panel) and initiate mitochondrial fragmentation and iRCD via Drpl and Bax recruitment, respectively.

CKM dysregulation is implicated in a wide variety of disorders and cancers (18). The finding that *Med13* ablation leads to early embryonic lethality (39) suggested that Med13 and Med13L have at least partially independent regulatory roles. However, our analysis of cyclin C, Cdk8, Med13 and Med13L promoter occupancy found that the CKM generally moves as a unit both to and from promoters, although some differences were observed (Fig. 2). In addition, with the possible exception of the repressed *Jmy* locus, Med13 and Med13L were found at all promoters analyzed and behaved similarly with respect to promoter recruitment or removal. These results may suggest that, at least for this gene subset, Med13 and Med13L have similar functions regulating transcription. Therefore, another interpretation of the knockout results is that removing Med13 or Med13L may invoke a gene dosage-related phenotype. For example, recent studies found that *MED13* or *MED13L* haploinsufficiency results in several overlapping, but not identical, developmental syndromes (40–45). Therefore, transcriptional control by Med13 and Med13L may be exquisitely sensitive to gene copy number. Eliminating both copies of *Med13* or *Med13L* may be sufficient to disrupt transcriptional control during embryogenesis sufficient to cause lethality. Analyzing the fate of mice heterozygous for both *Med13* and *Med13L* may address this question.

Our previous results from yeast revealed that Med13 destruction is required for the nuclear release of cyclin C following oxidative stress (27). These results fit well with structural studies indicating that Med13 tethers the CKM to the Mediator (17). Additional yeast studies found that the CKM is also disrupted in response to many stressors including starvation (46). However, closer examination revealed the fate of the CKM components changed depending on the stress. For example, in response to oxidative stress, the yeast Med13 is destroyed by the ubiquitin proteasome system (UPS) allowing cyclin C release into the cytoplasm to stimulate mitochondrial fragmentation and RCD prior to its destruction (28). In response to nitrogen starvation, Med13 is still destroyed but cyclin C has a different fate. Rather than undergoing nuclear release, cyclin C is degraded before cytoplasmic relocalization protecting the cell from its mitochondrial role promoting regulated cell death (47). Similar to yeast results, we found that in response to H_2_O_2_ or Torin1 treatment, cyclin C is removed from promoters that it represses under normal growth conditions (see Fig. 3B). However, several differences were observed. First, cyclin C levels do not change significantly following H_2_O_2_ or starvation stress. In addition, unlike yeast where the CKM is completely dismantled, this complex appears to remain mostly intact as it moves from one promoter to another. These differences may reflect the additional responsibilities the CKM has in mammalian cells. In addition to stress or developmental responsive genes, RNA-seq analysis also revealed a role for cyclin C in the transcriptional activation of genes involved in energy production and proliferation (7). Therefore, metazoans may have leveraged the requirement for cyclin C nuclear release with the increased role of the CKM in stress-independent transcription.

ChIP analysis revealed that CKM regulation was not always uniform. For example, H_2_O_2_ treatment resulted in nearly complete removal of cyclin C at the *Prkaa2* and *Crebrf* promoters, but only partial reduction in Cdk8, Med13 or Med13L. This pattern was not observed for other repressed genes, *Jmy* and *Gtf2h1.* These results suggest that the CKM was dissolved at the *Prkaa2* and *Crebrf* promoters. A potential explanation for these results is based on our finding of the mitochondrial role cyclin C mentioned above. In the cytoplasm, cyclin C mediates mitochondrial fragmentation and programmed cell death in MEF cells through its interaction with mitochondrial fission proteins Drp1 and Bax, respectively (25,26,48). These observations may indicate that the CKM at the *Prkaa2* and *Crebrf* promoters is disrupted to allow cyclin C nuclear release. In yeast, this release is triggered by Med13 destruction via the UPS. If a similar strategy is employed in MEFs, we would predict that MG-132 treatment would prevent CKM promoter release. In addition, failure to release the CKM would perhaps allow retention of its repressor function resulting in reduced transcription. These predictions were observed, albeit partially, with *Prkaa2* in which cyclin C was retained and reduced H_2_O_2_-induced activation was observed following MG-132 treatment (Fig. 4). However, not all repressed genes behaved in this manner and even at the *Prkaa2* promoter cyclin C still exhibited partial removal. These findings suggest that cyclin C release from promoters is facilitated by more than one mechanism that may be governed by locus-specific cues. Even at promoters that exhibit MG-132 sensitivity, there may be heterogeneity in the response. These findings suggest that the rules governing CKM movement and integrity are complex and may involve an intersection of signal transduction pathways, gene specific transcription factors and promoter chromatin contexts.

Our results are consistent with current models of the positive role for the CKM through one or more reported targets of Cdk8, including histone H3 or Cdk9 (P-TEFb) (46,49). The differences observed in the requirement of cyclin C for promoter activation may be due to the presence of locus-specific trans-activators. For example, the p53-dependent response network only partly requires cyclin C for full gene activation in H_2_O_2_-stressed cells. Conversely, *Adrm1* and *Gabarap* require cyclin C for both steady state and stress-induced transcription. Many of the genes induced by cyclin C following H_2_O_2_ treatment mediate iRCD. Together with the finding that Cdk8 kinase activity is required for transactivation, it is not surprising that Senexin A significantly reduced iRCD in response to oxidative stress. Interestingly, deleting *Ccnc* resulted in a more substantial reduction in iRCD efficiency in response to H_2_O_2_-induced oxidative stress treatment (Fig. 5D). The difference between the iRCD impact of deleting *Ccnc* and inhibiting Cdk8 may be due to the two, separate contributions of cyclin C. Similar to Senexin A treatment, *Ccnc* ablation inactivates Cdk8. However, the additional cytoplasmic role of cyclin C at the mitochondria may represent the additional impact on iRCD efficiency. Therefore, Senexin A treatment may demonstrate the transcriptional aspect of cyclin C-Cdk8-dependent iRCD. We have previously demonstrated that cyclin C nuclear release is not dependent on Cdk8 activity (24). Therefore, deleting *Ccnc* removes both functions resulting in more cellular protection from H_2_O_2_ treatment. This conclusion may be a bit simplistic as deleting *Ccnc* also removes the repressor function of the CKM. The contribution of the repressor aspect of cyclin C function to iRCD efficiency has yet to be separated from its other activities. Taken together, these findings reveal the contribution of cyclin C at multiple levels of the oxidative stress response pathway.

### Experimental Procedures

#### Cell Culture

*Ccnc^+/+^* (WT) and *Ccnc^-/-^* (24) immortalized MEF cells were cultured in Dulbecco’s Modified Eagle Medium (DMEM) supplemented with 10% fetal bovine serum (FBS) and 1% penicillin/streptomycin at 37°C and 5% CO_2_. WT and *Ccnc^-/-^* MEF cultures were grown to 75% confluence and treated as follows. Oxidative stress included exposure to 0.4mM H_2_O_2_ (Sigma Aldrich-CAS:7722-84-1) in DMEM without FBS for 4 h. Cells were treated with 250 nM Torin1 (Santa Cruz Biotech-CAS: 1222998-36-8) for 4 h. Cells were treated with 1 μM Senexin A (Tocris-CAS:1780390-76-2) for 24 h. Cells were treated with 5 μM MG-132 (Calbiochem-CAS:133407-82-6) for 1 h before oxidative stress treatment. The treatment protocols above were completed for 24 h for iRCD assay analysis. All experiments were conducted with three independent experiments conducted in triplicate for each condition.

#### Immunocytochemistry/Immunofluorescence (ICC/IF)

MEFs were cultured as described above on coverslips and then fixed with 4% paraformaldehyde for 10 min, permeabilized with 0.2% Triton X-100 for 10 min, blocked with 2% BSA, and incubated with primary antibody (1:2000 dilution of anti-cyclin C-Bethyl-A301-989A). Following a washing step, coverslips were incubated with secondary antibody (1:2000 dilution of anti-Rabbit Alexafluor^488^-Invitrogen-A-11008) and washed again before coverslips were mounted with 4’,6-diamidino-2-phenylindole (DAPI)–containing medium (Vector Labs). The images were acquired with a Nikon Eclipse 90i microscope equipped with a Retiga Exi charge-coupled device (CCD) camera and NIS software for data analysis. Mitochondrial staining was completed using MitoTracker Red CMXRos 30 min before fixing, as described by the manufacturer (Molecular Probes). Quantitation was accomplished with three independent cultures, 200 cells counted per sample. All microscopy experiments were conducted with three independent experiments conducted in triplicate for each condition.

#### Western Blot Analysis

Total cell lysate was prepared from cultured cells, as described above, in RIPA buffer (150 mM NaCl, 1% Igepal CA-630, 0.5% deoxycholate, 0.1% SDS, 50 mM Tris-HCl at pH 8, 5 mM EDTA, 20 mM NaF, 0.2 mM sodium orthovanadate and Sigma mammalian protease inhibitors), for 30 min with mild agitation at 4°C. Lysates were centrifuged at 13,000 × g for 10 min at 4°C. Protein concentrations were determined by the Bradford protein assay (Bio-Rad) and 30 μg of total protein was used for analysis. Western blot analysis for cyclin C was conducted with rabbit polyclonal primary antibody (1:2500 dilution) directed against cyclin C (Bethyl-A301-989A) and anti-rabbit secondary antibody (1:5000 dilution) (Abcam-ab6702). Western blot analysis for Med13 and Med13L were conducted with mouse monoclonal antibody (1:2500 dilution) directed against Med13 (Santa Cruz Biotech-sc515557) and anti-mouse secondary antibody (1:5000 dilution) (Abcam-ab6708); and rabbit polyclonal primary antibody (1:2500 dilution) directed against Med13L (Bethyl-A302-421A) and anti-rabbit secondary antibody (1:5000 dilution) (Abcam-ab6702). Western blots were incubated with CDP-STAR (CDP*) substrate and signals were visualized using an iBright FL1500 Imaging System (Thermo). β-actin (1:2500 dilution) (Abcam-ab8227) was used as loading controls. All western blot analyses were conducted with three independent experiments conducted in triplicate for each condition. The quantitation of Med13 and Med13L protein signals were completed using iBright Analysis software.

#### Flow Cytometry intrinsic Regulated Cell Death (iRCD) Assays

MEFs were seeded in culture conditions described above in 12-well plates at a density of 5 x 10^4^ cells and grown for 24 h_;_ before stress application for 24 h, as described above. Annexin V assays were conducted following the manufacturer’s instructions (BD Biosciences), using a BD Accuri™ C6 flow cytometer (BD Biosciences). All iRCD experiments were conducted with three independent experiments conducted in triplicate for each condition.

#### Chromatin Immunoprecipitation (ChIP) Assays

WT and *Ccnc^-/-^* MEF cultures treated as described above were crosslinked with 1% formaldehyde for 10 min at room temperature. The crosslinking reaction was quenched by the addition of 125 mM final concentration glycine. Cells were washed with cold PBS (containing Sigma mammalian protease inhibitors) then lysed with RIPA buffer (150 mM NaCl, 1% Igepal CA-630, 0.5% deoxycholate, 0.1% SDS, 50 mM Tris-HCl at pH 8, 5 mM EDTA, 20 mM NaF, 0.2 mM sodium orthovanadate and Sigma mammalian protease inhibitors), and subjected to sonication to produce DNA fragments < 500 bp. For immunoprecipitation, 1μg of antibody (α-cyclin C (Bethyl-A301-989A), α-Cdk8 (Santa Cruz Biotech-sc1521), α-Med13 (Santa Cruz Biotech-sc515557), α-Med13L (Bethyl-A302-421A) or α-GFP (Takara Bio-632375)), was added to samples and incubated for 8 h at 4 °C. The α-GFP antibody served as a non-specific control. Lysates were then incubated with magnetic Dynabeads (protein G) (Invitrogen-10004D) to collect immunoprecipitated samples. Following incubation, the beads were washed twice with RIPA buffer, four times with ChIP Wash Buffer (100 mM Tris-HCl pH 8.5, 500 mM LiCl, 1% [v/v] Nonidet P-40, 1% [w/v] deoxycholic acid), twice with RIPA buffer again, and twice with 1×TE (10 mM Tris-HCl, pH 8.0, 1 mM EDTA). Immunocomplexes were eluted for 10 min at 65°C with 1% SDS, and crosslinking was reversed by adding NaCl to a final concentration of 200 mM and incubating for 5 h at 65°C. DNA was purified for each sample, and a fraction (1/100) was used in qPCR amplification using ThermoFisher™ PowerSYBR™ Green PCR Master Mix and a StepOne™ Real Time PCR System. Primers specific to each individual gene promoter region were designed for quantification of the signal via the percent input method. Primer sequences for ChIP qPCR analysis can be found in Table S1. ChIP assays were conducted with three independent preparations in triplicate for statistical analysis via Student’s T-test.

#### RT-qPCR Analysis

Total RNA was prepared from the cell samples using Monarch® Total RNA Miniprep Kit. On-column DNase treatment was performed to eliminate contaminating DNA during RNA extraction. Total RNA (500ng) was converted to cDNA using the ThermoFisher™ Maxima cDNA Synthesis kit. The cDNA from each sample (1/100 dilution) was subjected to qPCR amplification using ThermoFisher™ PowerSYBR™ Green PCR Master Mix and a StepOne™ Real Time PCR System. These assays were conducted with three independent preparations assayed in duplicate. *Gapdh* was used as the internal standard for comparative (ΔΔC_T_) quantitation (50). Statistical significance was determined via Student’s T-test analysis. Primer sequences for RT-qPCR analysis can be found in Table S1.

## Acknowledgements

We thank Michael Law and David Stillman for helpful discussions and comments on this manuscript.

## Funding and other information

This work was supported by grants from the National Institutes of Health awarded to K.F.C. (GM113196) and R.S. (GM113052). Additional support was provided by the W.W. Smith Foundation (to K.F.C.) and the New Jersey Camden Health Initiative (to R.S.).

## Conflict of Interest

The authors declare there are not competing interests.

## Abbreviations

CKM: Cdk8 kinase module
mTOR: mammalian Target of Rapamycin
RCD: regulated cell death
iRCD: intrinsic regulated cell death
MEF: mouse embryonic fibroblast
ChIP: chromatin immunoprecipitation
UPS: ubiquitin-proteasome system

---

**Supplemental Table 1:**
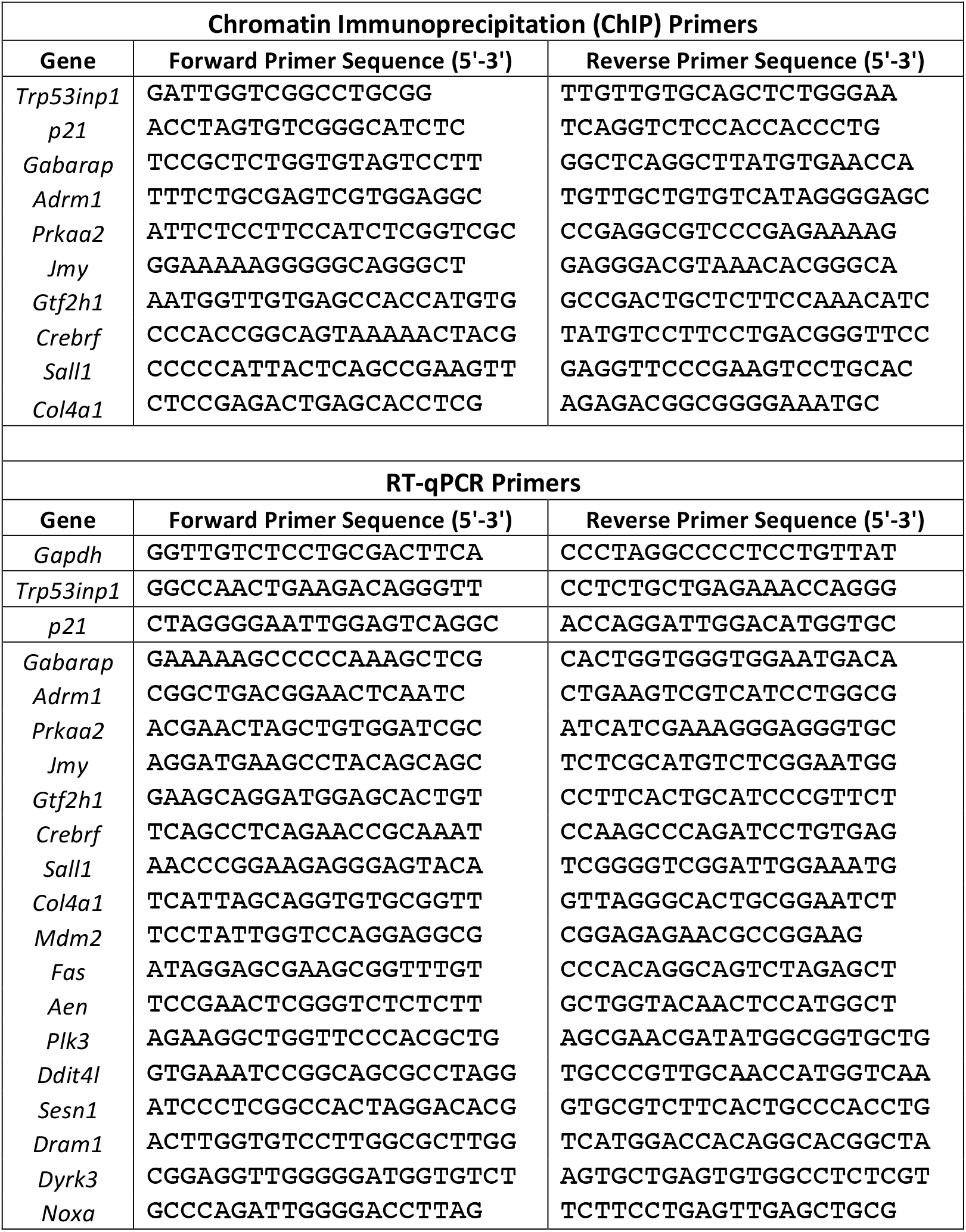
Primer sequences for ChIP and RT-qPCR studies.

**Figure SI:**
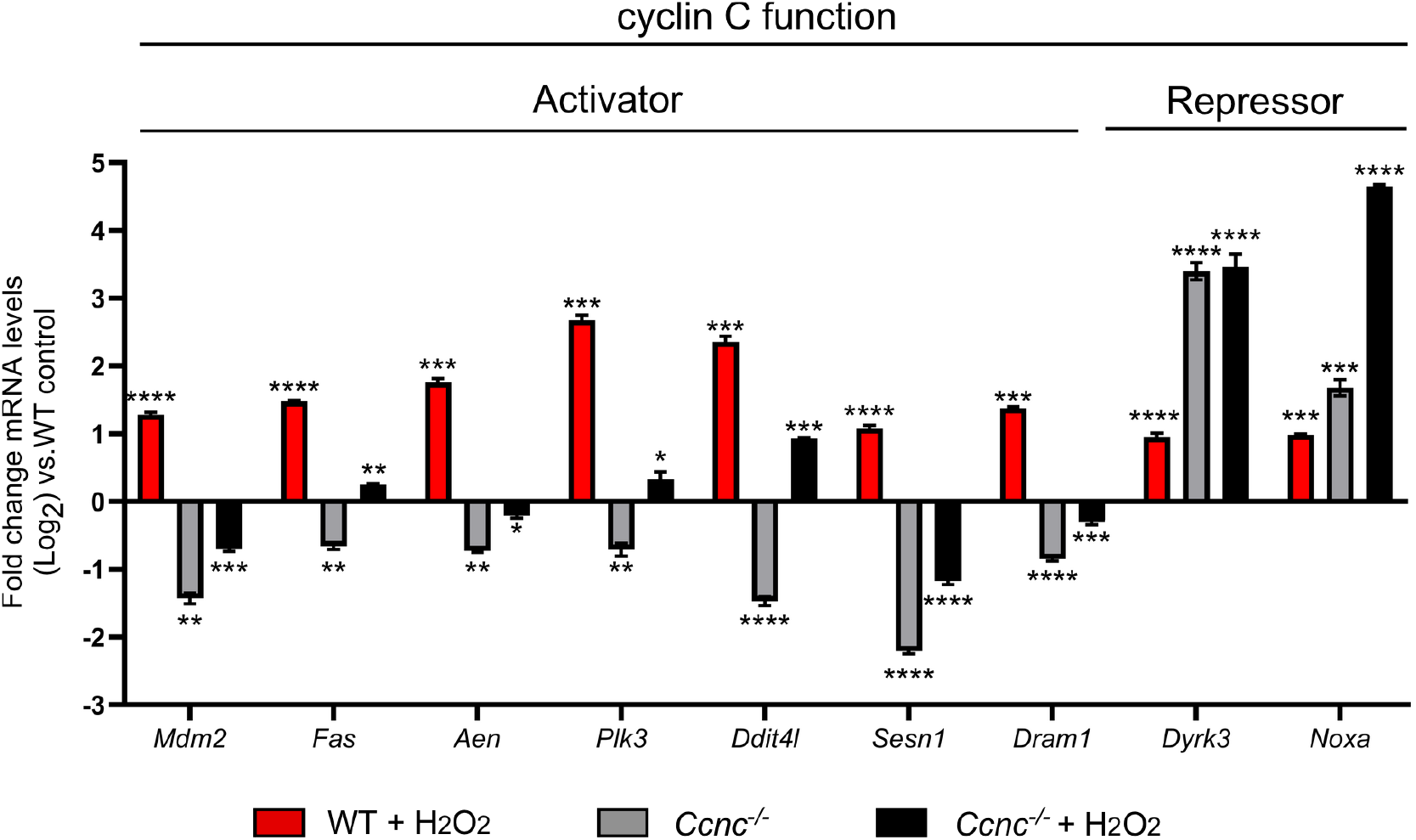
Cyclin C activator and repressor roles for H_2_O_2_-induced genes requiring p53. H_2_O_2_-induced genes identified by RNA-seq analysis were sorted into a cohort with the GO term “p53 regulated”. RT-qPCR analysis was performed on nine randomly selected genes in WT and *Ccnc^-/-^* MEF cultures before and after H_2_O_2_ treatment. Values shown are based on untreated WT control. Statistical significance is indicated by the following: * p-value < 0.05, ** p-value < 0.01, *** p-value < 0.001, **** p-value < 0.0001.

**Figure S2:**
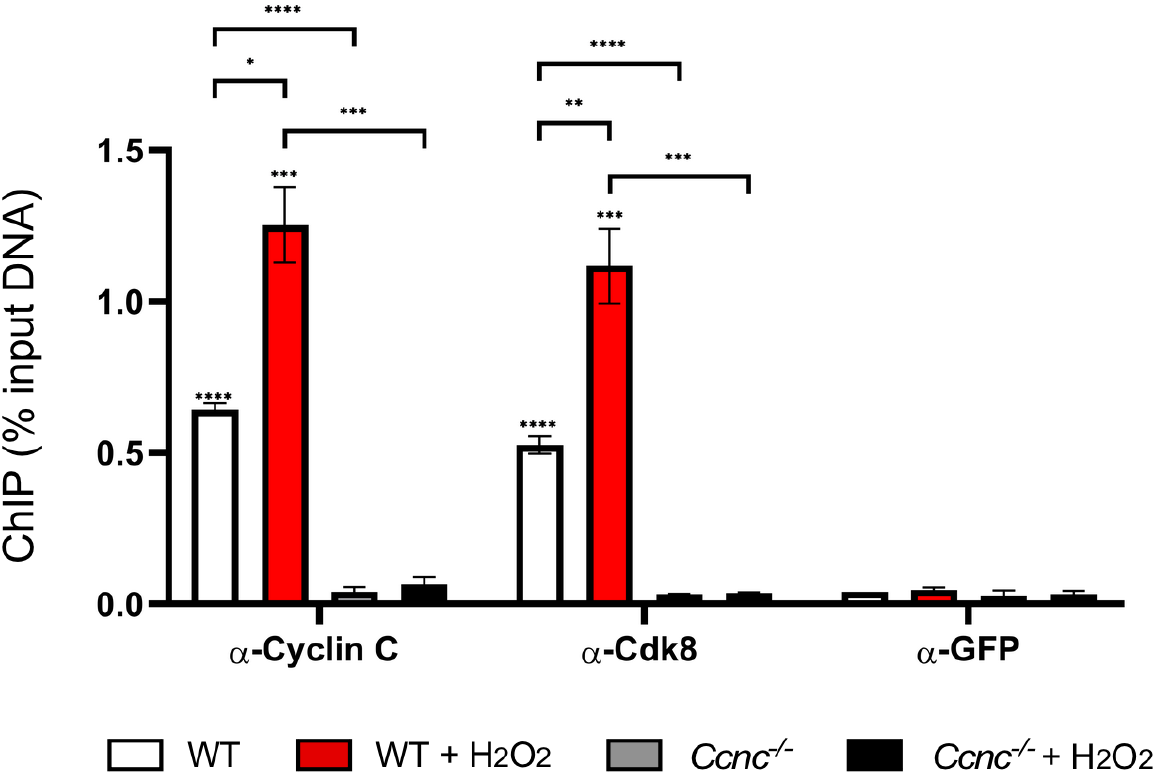
Cdk8 and cyclin C antibody specificity for ChIP experiments. ChIP analysis of *Trp53inp1* in the cell cultures, conditions and antibodies as indicated. Data obtained from RT-qPCR analysis from DNA purified from ChIP was calculated as percent input DNA. Statistical significance is indicated by the following: * p-value < 0.05, ** p-value < 0.01, *** p-value < 0.001, **** p-value < 0.0001

**Figure S3:**
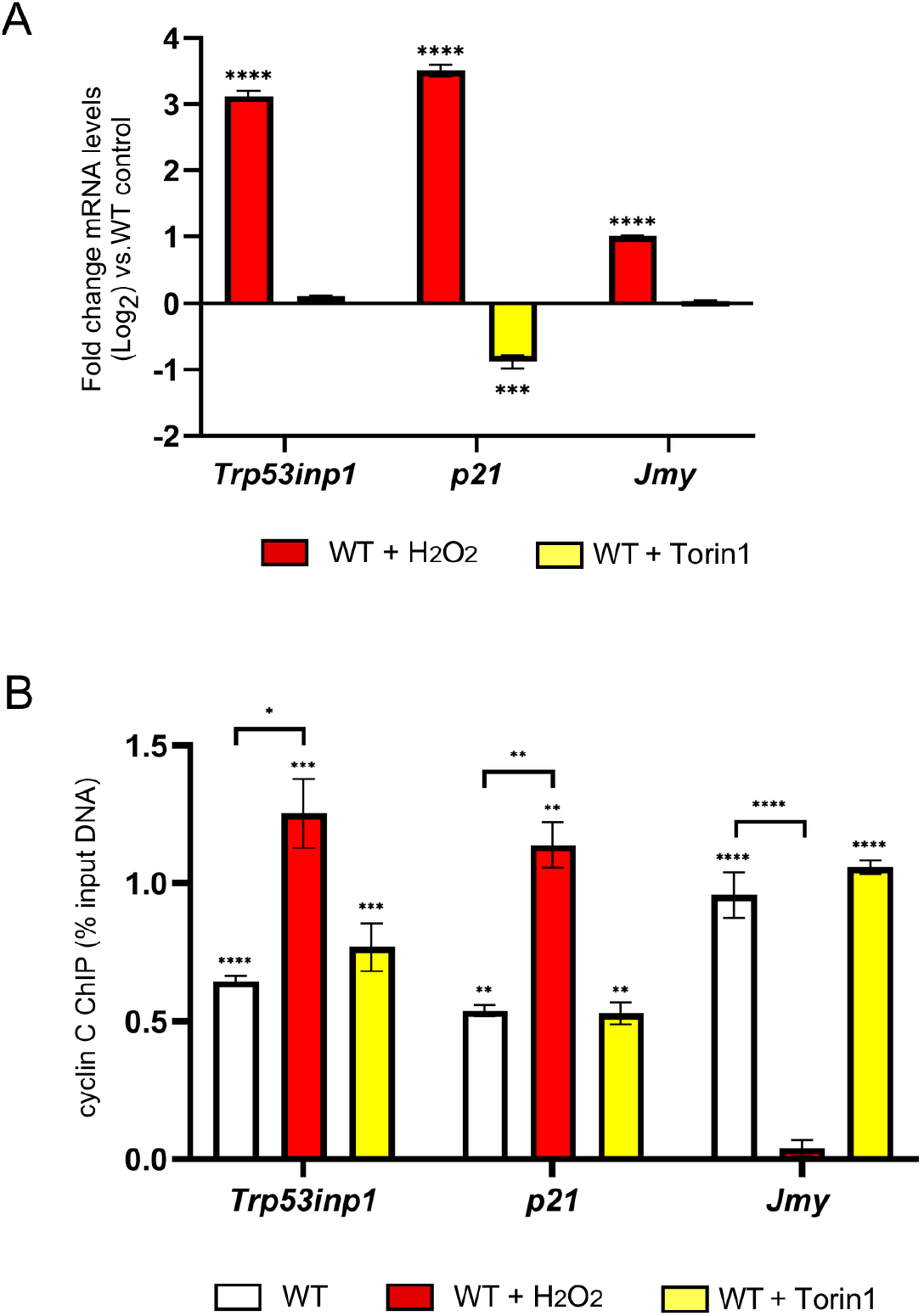
Genes within the p53-mediated response network are not induced during Torinl stress. RT-qPCR (A) and ChIP (B) data analysis on the genes indicated. Data obtained for mRNA levels and cyclin C-dependent ChIP were compared to an untreated WT control and a non-specific antibody (GFP), resepctively. Statistical significance is indicated by the following: * p-value < 0.05, ** p-value < 0.01, *** p-value < 0.001, **** p-value <0.0001.

